# Forest production efficiency increases with growth temperature

**DOI:** 10.1101/2020.04.15.042275

**Authors:** A. Collalti, A. Ibrom, A. Stockmarr, A. Cescatti, R. Alkama, M. Fernández-Martínez, G. Matteucci, S. Sitch, P. Friedlingstein, P. Ciais, D.S. Goll, J.E.M.S. Nabel, J. Pongratz, A. Arneth, V. Haverd, I.C. Prentice

## Abstract

We present a global analysis of the relationship of forest production efficiency (FPE) to stand age and climate, based on a large compilation of data on gross primary production and either biomass production or net primary production. FPE is important for both forest production and atmospheric carbon dioxide uptake. Earlier findings – FPE declining with age – are supported by this analysis. However, FPE also increases with absolute latitude, precipitation and (all else equal) with temperature. The temperature effect is opposite to what would be expected based on the short-term physiological response of respiration rates to temperature. It implies top-down regulation of forest carbon loss, perhaps reflecting the higher carbon costs of nutrient acquisition in colder climates. Current ecosystem models do not reproduce this phenomenon. They consistently predict lower FPE in warmer climates, and are therefore likely to overestimate carbon losses in a warming climate.

## Introduction

Autotrophic respiration releases to the atmosphere about half (~60 Pg C yr^−1^) of the carbon fixed annually by photosynthesis^1^. Forests are the largest carbon sink on land, taking up about 3.5 ±1.0 Pg C yr^−1^ (2008–2017) on average^2^. A small change in the proportion of carbon losses, for example due to climate change, would thus strongly affect the net carbon balance of the biosphere. Predicting the carbon balance of forests under changing climate requires understanding of how much atmospheric CO2 is assimilated through photosynthesis (gross primary production, GPP), how much is released due to plant metabolism (autotrophic respiration, *R*_a_), how efficiently plants use assimilated carbon for the production of organic matter (net primary production, NPP), and how organic carbon is partitioned into plant organs (biomass production, BP) versus other less stable forms – which include soluble organic compounds exuded to the rhizosphere or stored as reserves, and biogenic volatile organic compounds (BVOCs) emitted to the atmosphere.

The climate sensitivity of the terrestrial carbon cycle can be benchmarked using ratios between these fluxes across a range of climates. We focus here on the ratio of NPP to GPP, the called ‘carbon use efficiency’ (CUE) and of BP to GPP, called ‘biomass production efficiency’ (BPE). The two concepts are inherently close, but not identical^3,4^. BPE is substantially easier to obtain, however, because the additional fluxes that constitute NPP are notoriously difficult to measure. For this reason there are far more data available on BPE, while uncertainties associated with both BP and NPP measurement make it impossible to distinguish them in large data compilations.

Over twenty years ago, the debate about spatial gradients of forest CUE seemed to be resolved by Waring et al.^5^, who found CUE to be nearly constant (0.47 ±0.04 s.d.: here and elsewhere, ± denotes one standard deviation) across a range of temperate and boreal forest stands (*n* = 12). The assumption of a universal value for CUE – implying a tight coupling of whole-plant respiration to photosynthesis – has obvious practical convenience, and numerous vegetation models have adopted it^4^. Complex process-based vegetation models however assume decoupling of photosynthesis and respiration, with the latter driven by temperature^6^ and biomass^7^ – implying that CUE must vary. There is no general, observationally based consensus as to which of these two (mutually incompatible) model assumptions is nearer to the truth. One study found that BPE is greater at higher soil fertility^3^, perhaps because less carbon investment is required for nutrient acquisition. Forest management^8^, stand age^9^ and climate^10,11^ have also been implicated as influences on CUE and BPE.

Here we revisit the global patterns of forest CUE and BPE allowing both for the possibility of multiple controls, and for the potential effects of methodological uncertainty, based on an unprecedentedly large global set of data on forest CUE and/or BPE (*n* = 244) spanning environments ranging from the tropical lowland to high latitudes and high altitude (Figure S1).

## Results

Results show that both CUE (0.47 ±0.13; range 0.24 to 0.71; *n* = 47) and BPE (0.46 ±0.12; range 0.22 to 0.79; *n* = 197) have large spatial variability; therefore, neither can be assumed to be uniform (Figure 1a-b). CUE and BPE are statistically indistinguishable in our dataset because of uncertainties associated with both quantities (±0.39 for CUE and ±0.16 for BPE: see Methods). Therefore we assessed estimates of both BPE and CUE as a single metric, hereafter called ‘forest production efficiency’ (FPE). The average FPE in our dataset (0.46 ±0.12; range 0.22 to 0.79; *n* = 244) is statistically indistinguishable from that provided by Waring et al.^5^, but its standard deviation is three times larger (Methods and ref.^4^). Different GPP estimation methods (Methods) produced slightly different distributions (Figure 2a), with median values ranging from 0.44 (*scaling*) through 0.48 (*micrometeorological*) to 0.49 (*model*). Stand age had a further effect on the average FPE, as shown by the differing median CUE and BPE values of stands in intermediate (in the silvicultural sense, i.e. 20 to 60 years) (Figure 2b) and younger age classes. Figure 3 shows how the data compare to those published by Waring et al.^5^. The very small variability of CUE reported by Waring et al.^5^ was already noted by Medlyn & Dewar^12^ as untypical, and artificially constrained by the method used to calculate CUE. Medlyn & Dewar^12^ suggested a 0.31 to 0.59 range as being more realistic. Figure 3 also indicates systematically lower values than Waring et al.^5^ for forests with GPP < ~2,000 g C m^−2^ yr^−1^, especially in forests in the old age class; and a tendency to higher values for forests with GPP > ~2,000 g C m^−2^ yr^−1^ and in the young age class.

**Figure 1.**
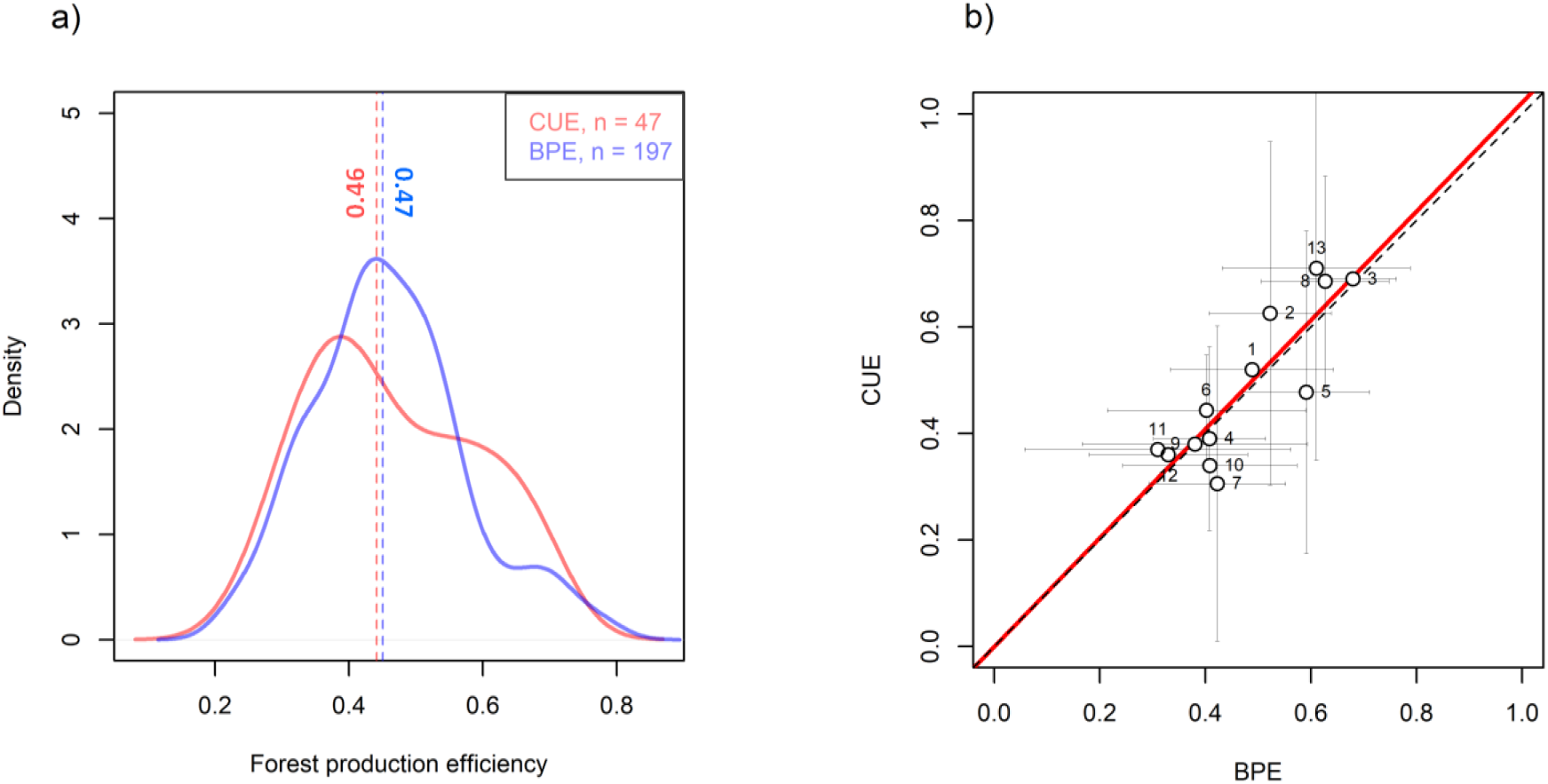
Carbon use efficiency vs. biomass production efficiency. **a**, Density plot of Carbon Use Efficiency (CUE, *n* = 47) and Biomass Production Efficiency (BPE, *n* = 197) data from all available data. The vertical lines are medians. **b**, Scatter plot and linear regression line of site data where both CUE and BPE were available, forced through the origin (adjusted R^2^ = 0.98, *slope* = 1.022 ±0.041 S.D.: i.e. not significantly different from 1).

**Figure 2.**
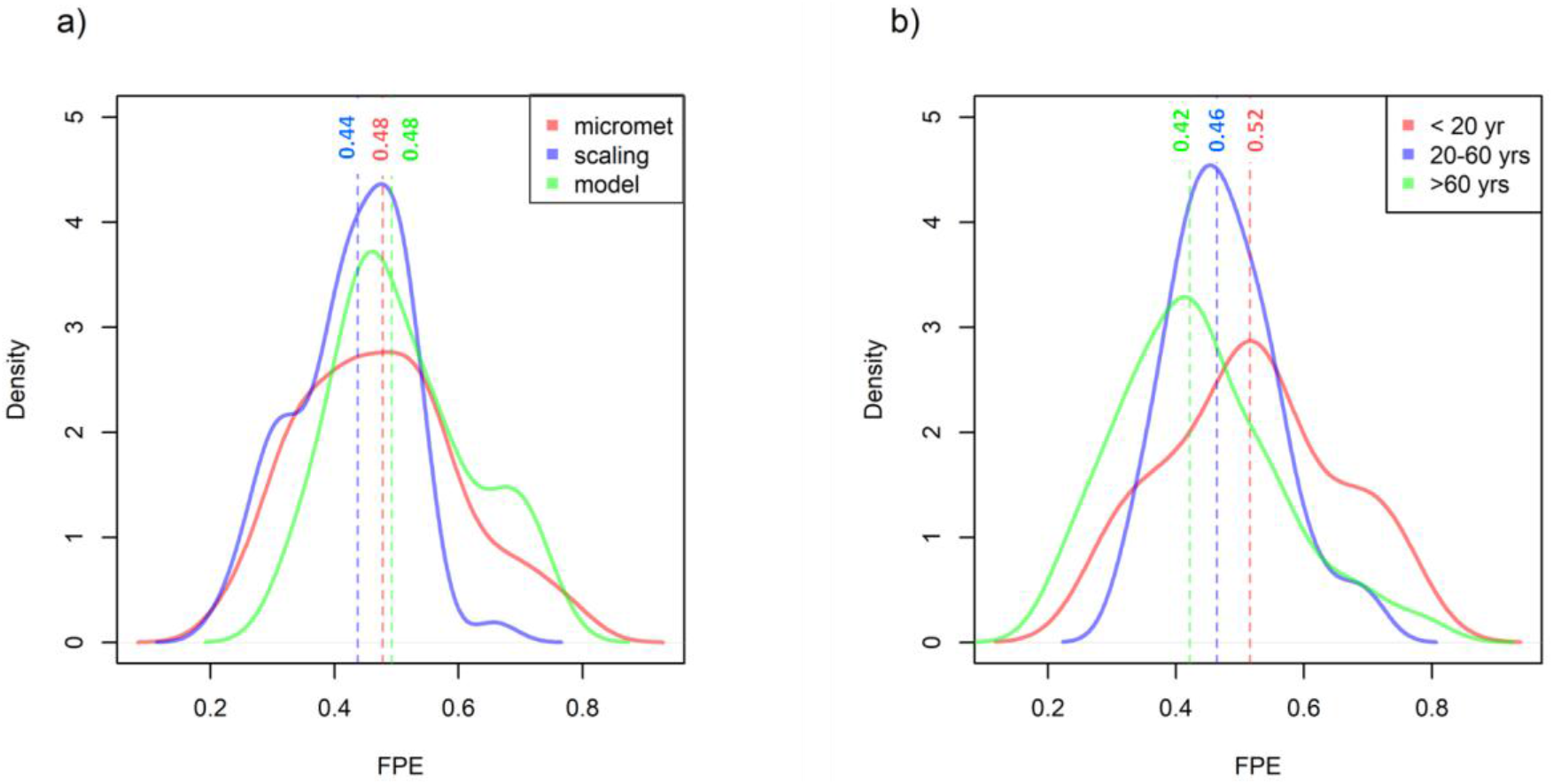
GPP method and age effects on FPE variability. **a**, Forest production efficiency density plots for three subsets of data where the GPP was estimated with three different methods (*n* = 98, 73 and 53 for micrometeorological, scaling and models, respectively). The vertical lines are medians. **b**, Density plots for different age classes (*n* = 47, 49 and 77, for age < 20, 20-60 and > 60 years, respectively).

**Figure 3.**
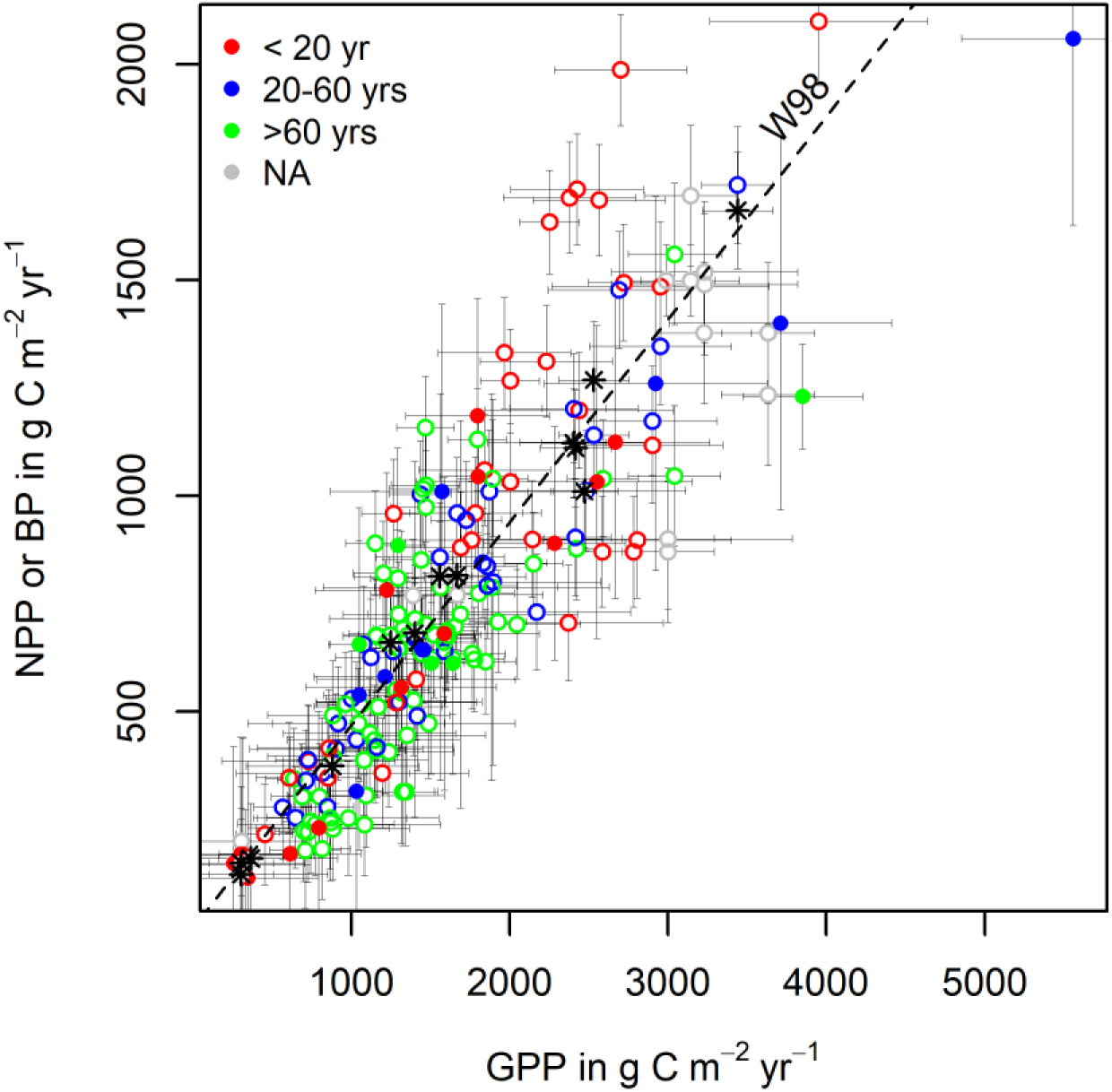
Current work dataset vs. Waring et al.’^5^ hypothesis. Scatter plot of net primary production (NPP, g C m^−2^ yr^−1^) or biomass production (BP, g C m^−2^ yr^−1^) versus gross primary production (GPP, g C m^−2^ yr^−1^). Open circles: BP, filled circles: NPP, stars from Waring et al.^5^. Age classes are marked by colours; NA stands for ‘age not available’. The uncertainty (g C m^−2^ yr^−1^) of the data points is indicated by bars (for data uncertainty see Methods). The line marked with W98 represents a CUE of 0.47, as reported by Waring et al.^5^.

We use mixed-effects multiple linear regression to infer the multiple drivers on spatial gradients of FPE. This method separates the contribution of every predictor variable included in the analysis, even if they are correlated to some degree (Methods). Four predictors – out of an initial selection of eleven (listed in the online Material, Methods) – proved to be important: stand age (age, yrs), mean annual temperature (MAT, °C), total annual precipitation (TAP, mm yr^−1^) and absolute latitude (|lat|, °), all included as a fixed effects. *GPPmethod* – the method used to measure GPP – was included as a random effect (Table 1, eq. (1)).

**Table 1.**
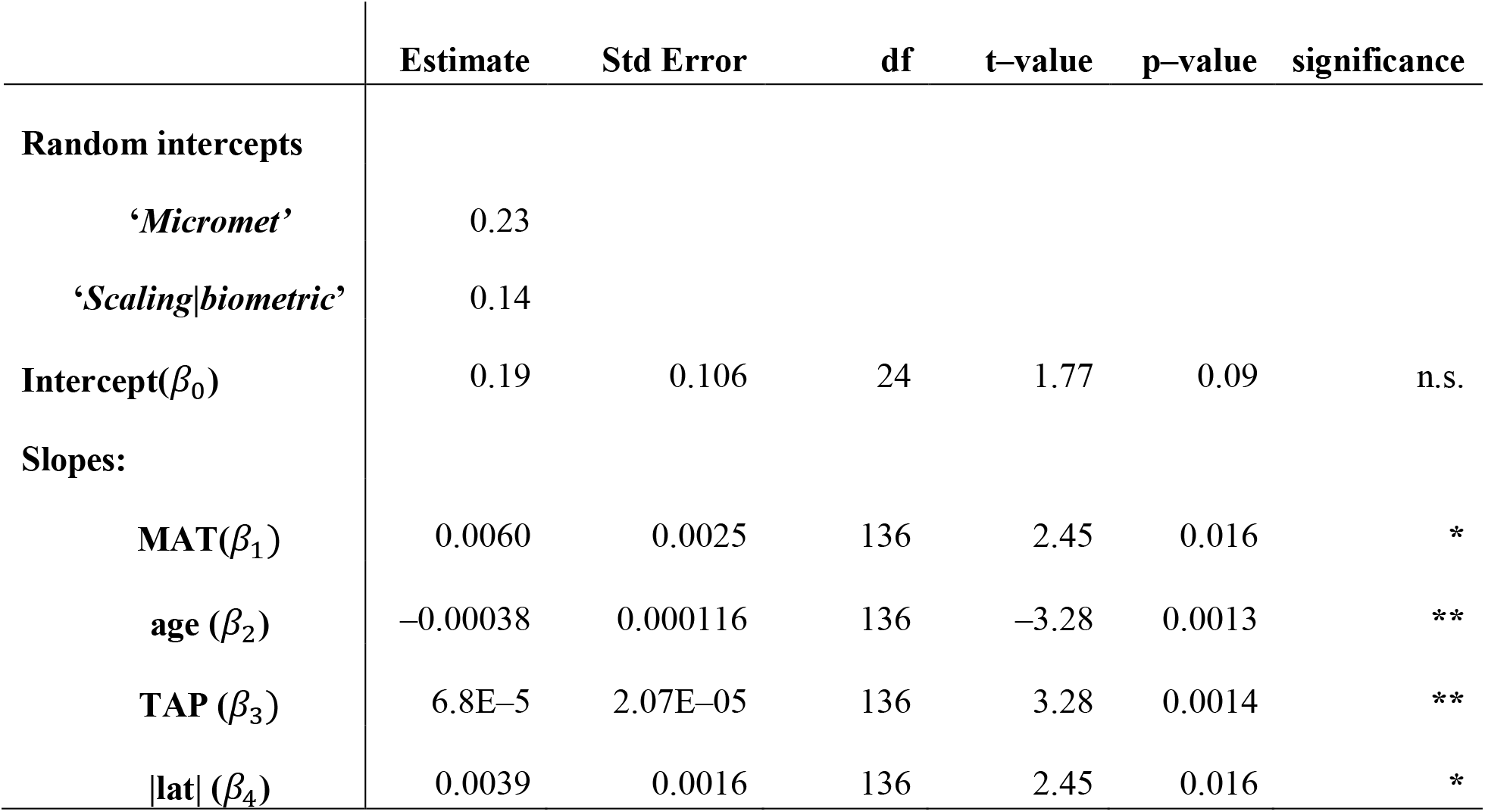
Parameters of the mixed-effects multiple regression model (equation 1). Parameter estimate of coefficients in equation (1) and their standard errors (Std. Error), degrees of freedoms (df), t- and p- values of the T test and the ANOVA(* p < 0.05, ** p < 0.01, *** p < 0.001). The squared Pearson’s correlation coefficient value is 0.3.

The use of multiple regression was essential for this analysis. Simple correlations between FPE and individual predictors showed no significant effects, while there were large and significant correlations between the different predictors (Table S1). The model with four fixed effects (equation 1) could not be further reduced at a 5% test level, i.e. omitting any one of these predictors yielded a model that significantly differed from the model with all parameters (Table S2 lists the combinations that were tried):

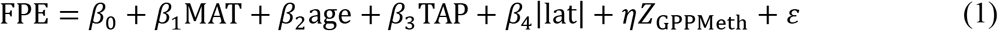

where *β*_0_ is the intercept, and *β*_1_-*β*_4_ are the estimated sensitivities of FPE to MAT, stand age, TAP and |lat|; *ηZ*_GPPMeth_ is a random intercept for the two *GPPmethod* classes and *ε* is the residual (Table 1). We examined random effects from three methods to estimate GPP (Methods). The methods *biometric* and *scaling* turned out to constitute a single class, while GPP values determined from *micrometeorological* measurements were systematically higher (thus FPE was systematically lower). The multiple regression model explained 30% of the variance in the observed FPE values. Given the large uncertainty in the estimation of NPP and GPP values, and the structural and physiological diversity of the forests, this value was unexpectedly high.

We could not fit an independent statistical model for CUE, because there were too few sites with NPP (*n* = 31) measurements. However, adding a random intercept for the two categories (CUE or BPE) to equation (1) yielded almost identical values, of 0.47 for CUE and 0.46 for BPE.

We also applied the mixed-effects multiple regression model to the TRENDY v.7 outputs of eight Dynamic Global Vegetations Models (DGVMs) to examine whether the multivariate relationships shown for the FPE data could also be seen in the model simulations, testing whether the observed pattern would emerge from physiological processes and allocation schemes as used in the models. We originally aimed to use the same mixed-effects linear model, at the same locations – i.e. where the observed data come from – to fit the simulated FPE. However, because these DGVMs do not consider forest age, we needed to adapt the model equation to:

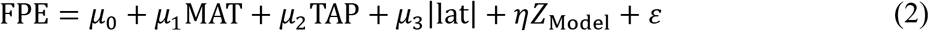

Neither this model, nor any of the models that could be derived from this equation, fulfilled the conditions of normally distributed residuals. In other words, the simulations did not represent a common emergent relationship consistent with the data. The most likely explanation is that the models use different parameters (and even sometimes different functional relationships) for different biomes, so that no general relationship applying across all forest types can be expected to emerge. This phenomenon is evident on inspection of Figure 5, which reveals that many models have discontinuities in CUE across the variable space.

**Figure 4.**
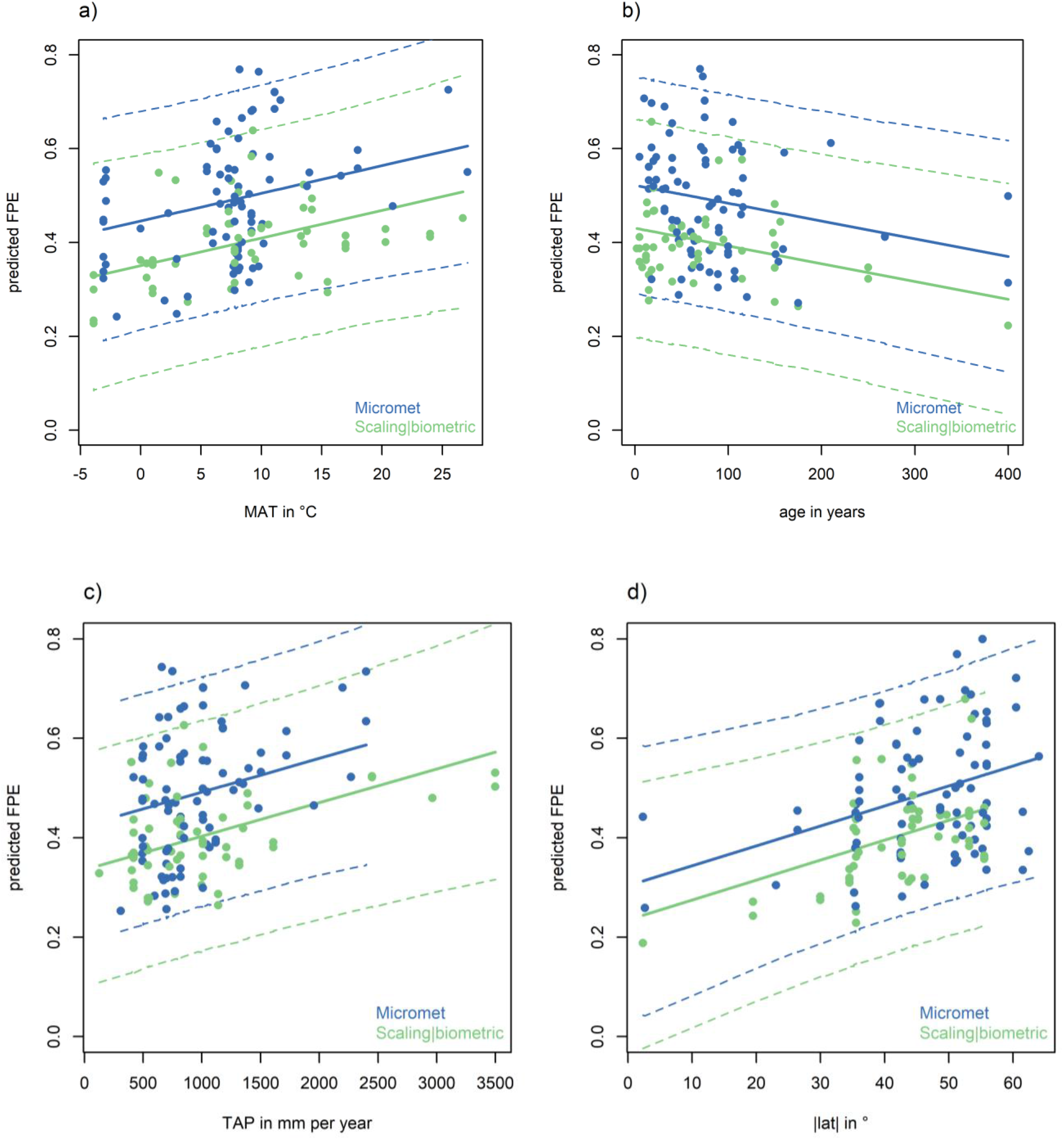
Predicted FPE vs. single effects of environmental and structural variables. Predictions of the mixed linear model for single fixed effects (equation 1), given the other independent variables constant at their average values for that GPP method category. The dashed lines represent confidence intervals at the 0.05 and 0.95 levels calculated with the function ‘predict Interval’ of the R-package `‘merTools’`.

**Figure 5.**
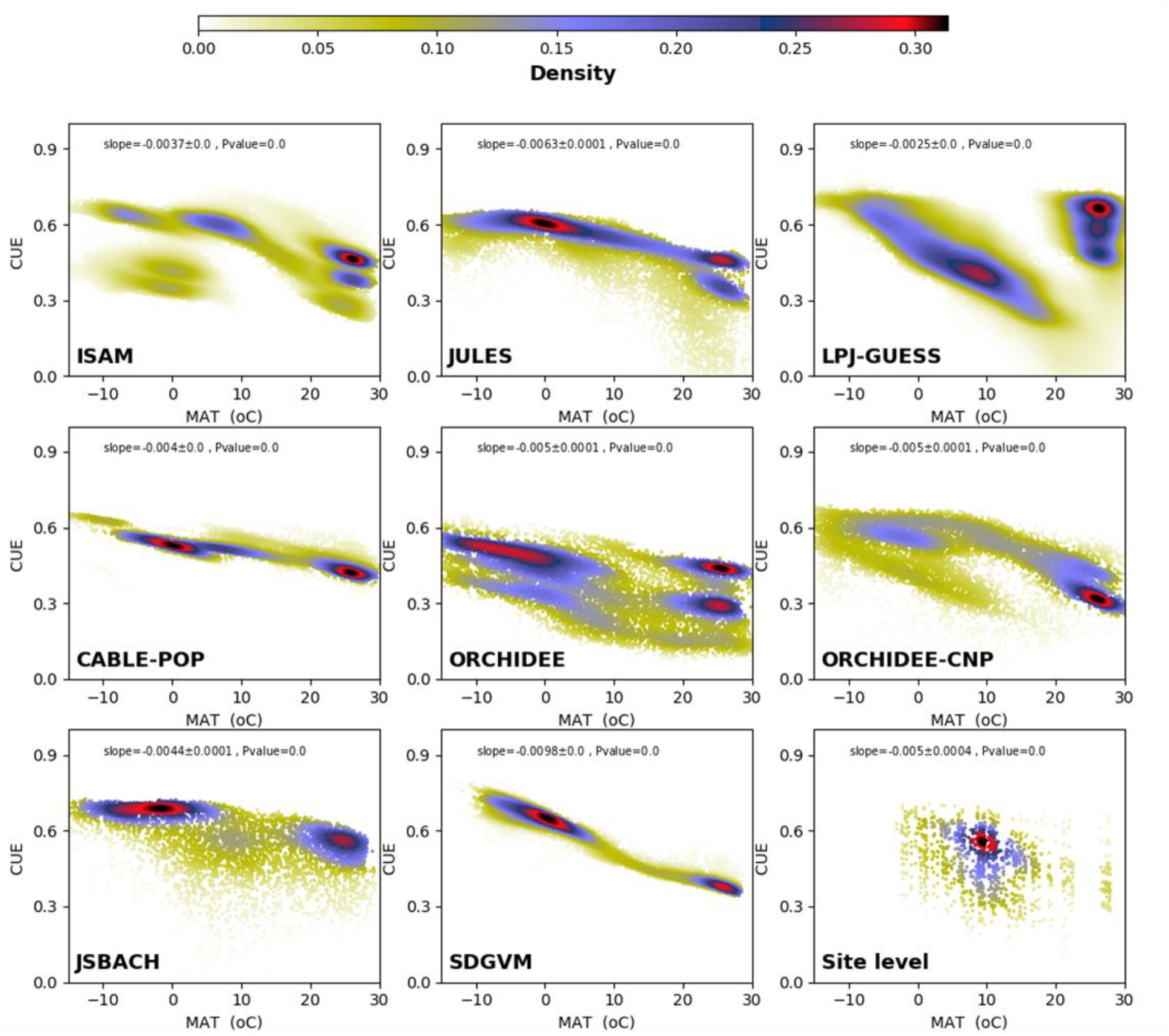
Modelled TRENDY v.7 CUE and growth temperature patterns. Density plots (i.e. frequency of a CUE value divided by the total number of grid cells) of simulated forest carbon use efficiency (CUE) derived from TRENDY v.7 process-based models: ISAM, JULES, LPJ-GUESS, CABLE-POP, ORCHIDEE, ORCHIDEE-CNP, JSBACH and SDGVM, averaged from 1995 to 2015 as function of MAT (°C). In the last (right bottom) density plot, data points extracted from coordinates of observed sites and used to plot the simulated CUE as function of MAT from the eight TRENDY v.7 models.

CUE outputs from the TRENDY v.7 model ensemble, produced by eight Dynamic Global Vegetation Models (DGVMs), consistently showed a negative relationship with MAT – opposite to that shown by our analysis of data. The slope (*d*CUE/*d*MAT) from observed data was +0.0015 °C^−1^; the slopes from models ranged from –0.0025 °C^−1^ for LPJ-GUESS to –0.0098 °C^−1^ for SDGVM (Figure 5). The average slope across the eight models was –0.005 °C^−1^. All models showed high CUE for boreal forests and low CUE for tropical forests, but with considerable variation among models (Figure S3). The modelled CUE values agree well with the data only in temperate regions (MAT 5 to 15 °C, *n* = 156), but differ greatly in boreal (MAT < 5 °C, *n* = 35) and tropical (MAT > 15 °C, *n* = 40) regions.

## Discussion

### The empirical ranges of CUE and BPE and the age-related effects on FPE

Under some extreme circumstances, the carbon flux to mycorrhizae and root exudates can constitute as much as 50% of daily assimilation^13^ or as much as 30% of annual NPP^14,15^. While in non-stressed conditions BVOCs consume a small fraction (~5% or less) of annual NPP, under stressed conditions and in hot climates, BVOC emissions can also consume from 15 to 50% of annual NPP^16,17^. Thus CUE – if not equal to BPE – should always be larger than BPE. In our data set, in those cases where both could be estimated, CUE was larger than the estimated BPE in seven out of thirteen cases (Supporting Information, Table S3). In the other cases the estimated CUE was statistically not distinguishably from BPE. This finding suggests that the fraction of these occult organic carbon flows varies substantially across different forests.

Statistically fitted values of BPE and CUE ranged between 0.27 (–0.04) and 0.58 (+0.04). The numbers in parentheses for CUE and BPE reflect our estimates of methodological bias (random intercepts, see Table 1). Ninety-two percent of BPE and CUE values in the dataset lie within the ‘allowable’ range reflecting maximum growth with minimum expenditure (0.65) and minimum growth with maximum maintenance costs (0.2) according to Amthor^18^. The remaining 8% of the data exceed Amthor’s upper bound. However no values below 0.2 were found, and it seems likely such values cannot be physiologically sustained by plants for long periods^7^. They might be encountered in moribund stands, unlikely to be sampled (perhaps an example of the ‘survivorship bias’ or ‘desk drawer problem’^19^). However, values > 0.65 apparently can temporarily occur in young, actively growing forests.

Age- or size-related declines in both GPP, NPP and FPE (as CUE or BPE) have been reported in earlier studies ^8,10,20,21^ but a decline in FPE is shown unequivocally here (slope *d*FPE/*d*age = –0.0004 yr^−1^), based on a substantially larger dataset than previously analysed^22–24^(Figure 4 and Table 1). This decline could have several contributory causes. First, the longer transport pathway for water in taller trees can result in more closed stomata (avoiding xylem cavitation) and therefore reduced GPP^25^, with no corresponding reduction in the short-term in *R*_a_. Second, larger trees may respire more because of their greater sapwood volume and mass per unit leaf area^7,26,27^, leading to increased *R*_a_ (for the maintenance of living sapwood tissues) and reduced NPP relative to GPP. Third, soil fertility declines due to nutrient immobilization as stands age^28^; this is consistent with observations of an increased ratio of fine-root-to-leaf-carbon, and reduced nitrogen concentration in soils^9,29^. Ontogenetic shifts from structural biomass to reserve allocation, and structural and resource limitations in older stands, are thus all expected to decrease production efficiency^30^. Conversely, reducing plant competition and rejuvenating stands through forest management should tend to increase both CUE and BPE^8,31^. At the stand scale, closing canopies may contribute further to reducing or stabilizing GPP^32^ at the level of individual trees. It is also likely that young trees allocate more carbon to biomass growth as they compete spatially for light and nutrients; while older trees invest more in maintenance of their existing biomass, and prioritize the chemical defence of that biomass, relative to acquisition of new biomass^33,34^.

An additional hypothesis^35^ invokes an increase in non-structural carbohydrates (NSC) allocation as trees grow. NSC is a substantial carbon pool, containing one to four times the carbon content of leaves in the canopy and increasing as trees increase in size^36^. An increased flux to NSC however would imply a reduced BPE, not a reduced CUE.

### Environmental effects on FPE

The observed increase in FPE with increasing TAP is rather new, to our knowledge, in literature. Higher TAP results in increasing soil water availability and greater stomatal openness which might imply increased photosynthesis. There is no direct evidence that water availability influences autotrophic respiration; on the other hand, respiration has been found to increase with drought^37^. With increasing TAP, photosynthesis might be expected to increase faster than plant respiration does, leading to higher CUE and BPE.

The increase of FPE with absolute latitude has not been described before. Higher latitudes cause longer days in summer. Diffuse radiation fractions also increase with the path-length of radiation through the atmosphere. Both effects might be expected to increase the radiation use efficiency of GPP. While high irradiance in the tropics leads to saturation of photosynthesis in the uppermost leaf layers^38^, they also allow for higher leaf area to utilize the transmitted radiation in the relatively short daylight hours. Higher leaf area also implies higher *R*_a_ and lower FPE. The latitude effect compensates for the MAT effect, because these two variables are negatively correlated. Despite the correlation, the mixed linear model is able to distinguish between the individual effects of MAT and |lat|. This can be demonstrated by comparison of high-latitude with high-elevation sites at the same MAT. The effects of radiation regimes were incorporated in models which, therefore, could in principle represent their effect on FPE.

The observed increase in FPE with MAT is new, and is opposite to what would be expected based on the instantaneous responses of photosynthesis and plant respiration as described in textbooks^39^ and assumed in many process-based models (Figure 5). The instantaneous response of respiration to a temperature change is steeper than that of photosynthesis^40^. Moreover, under natural conditions photosynthesis is commonly limited by light while respiration is not. However, the instantaneous response of autotrophic respiration rate is largely irrelevant here because of the longer time scale. A long line of investigations, starting with Gifford^41^, has shown the ubiquity of respiratory thermal acclimation, whereby the effect of increased growth temperature on enzyme kinetics is offset by a lowering of the base rate^42^. This acclimation takes place on a time scale of days to weeks^1^. Genetic adaptation throughout multiple generations is expected to proceed in the same direction (for definitions and distinctions between *acclimation* and *adaptation* see ref.^43^). One consequence of these processes is that observed rates of respiration vary with temperature (in both space and time) far less steeply than would be expected based on the instantaneous response of enzyme kinetics^44^. This has been shown comprehensively in leaves, and is likely to apply to all plant tissues^1^. Moreover, the ratio of respiration to carboxylation capacity, assessed at growth temperature, is slightly but significantly larger in colder climates^44^.

He et al.^45^, using MODIS-NPP product and the outputs from five process-based carbon cycle models, found – in contrast to our results – a strong latitudinal pattern with higher CUE at high latitudes declining non-linearly with increasing MAT and stabilizing at increasing TAP. He et al.’s^45^ results were obtained using an ‘emergent constraint’ method, which implicitly assumes that key relationships common to a set of models are correct. Thus, He et al.’s^45^ findings may ultimately reflect the standard assumption of models (including the MODIS-NPP product) that *R*_a_ increases with temperature more steeply than GPP^40^. We have shown the same modelled patterns here in the TRENDY v.7 ensemble. However, our analysis indicates that this assumption is incorrect, and shows that the same error is present in all the participating models.

Adaptive mechanisms, potentially contributing to respiratory thermal acclimation, include changes in the physiology and growth of active tissues (i.e. the relation between assimilating and non-assimilating tissues) and changes in the amount of enzymes and their activation states to match substrate availability^40,46^. Heat tolerance in leaves has also been found to increase linearly with temperature and latitude^47^. Therefore, a simple explanation for the increase of FPE with temperature is that plant can achieve the same function at a higher temperature with smaller amounts of enzymes, thereby decreasing the respiratory losses incurred during the maintenance of catalytic capacity. Especially low FPE in boreal forests could be the consequence of greater allocation of assimilates to nutrient acquisition (via root exudation and exports to mycorrhizae) in cold soils where microbial activity is much lower than in tropical forests^29^. Low FPE in cold climates may also reflect the need to repair tissues affected by frost damage^48^.

### Whole-plant constraints and consequences for modelling

Amthor^18^ derived an upper bound of 0.65 for CUE, based on a rough quantification of the minimum respiratory costs for plants to function. His lower bound of 0.2 was based on the need for a sufficiently positive carbon balance to have minimum photosynthesis to survive and to make trees able to compensate for tissue turnover, reproduction and mortality. However, most CUE values lie within narrower bounds, suggesting the existence of additional regulatory mechanisms at the whole-plant scale. Gifford^41^ noted that respiration and primary production are interdependent, because carbon must be assimilated before it is respired, while respiration is required for the growth and maintenance of tissues. He opined that: *‘Plant respiratory regulation is too complex for a mechanistic representation in current terrestrial productivity models for carbon accounting and global change research’* and indicated a preference for simpler approaches that capture the ‘essence’ of the process. The opposite view was expressed by Thornley^49^, who argued that: ‘*attempting to grasp and pin down complexity is often the first step to finding a way through a labyrinth*’. Without need to necessarily take a position on this controversy, we note that the standard approach in today’s land ecosystem models as shown here, or more generally in vegetation models – where autotrophic respiration per unit of respiring tissue is typically determined as a fixed basal rate at a standard temperature (commonly 15 or 20 C°), increasing with the substrate and temperature according to a fixed Q_10_ factor or Arrhenius-type equation – cannot generate the observed positive response of CUE or BPE to growth temperature. Moreover, as shown in Figure 5, the presence of discontinuities in CUE probably represents an attempt to sidestep an inevitable consequence of this incorrect approach: unless plant functional types from warmer environments are assigned lower basal respiration rates, modelled CUE becomes implausibly low in warm climates. However, the idea of assigning fixed basal respiration rates to plant types has no observational or experimental basis.

In contrast, the use of production efficiency concepts in models seems well motivated^50^, provided they are not assumed to be constant across different stands and environments. Production efficiency is a valuable unifying concept for the analysis of forest carbon budgets. Although more variable than was once thought, FPE appears to be a relatively conservative quantity, subject to inherent biological constraints, that has demonstrable relationships to stand development, latitude and climate. The possible explanations for the observed global multi-factorial pattern in FPE give rise to hypotheses on how vegetation models might incorporate whole plant regulation mechanisms of the carbon losses for a given stand. The demonstrated empirical pattern should then be used to constrain new model developments.

## Methods

### Definitions of terms

GPP is defined here as ‘the sum of gross carbon fixation (carboxylation minus photorespiration) by autotrophic carbon-fixing tissues per unit area and time’^51^. GPP is expressed as mass of carbon produced per unit area and time, over at least one entire year. NPP consists of all organic carbon that is fixed, but not respired over a given time period^51^:

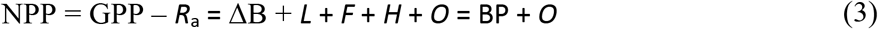

with all terms expressed in unit of mass of carbon per unit area and time. *R*_a_ is autotrophic respiration; the annual change in standing biomass carbon is ΔB; litter production (roots, leaves and woody debris) is *L*; fruit production is *F*; the loss to herbivores is *H*. BP is biomass production^3^. Symbol *O* represents ‘occult’ carbon flows, i.e., all other allocations of assimilated carbon, including changes in the non-structural carbohydrate pool, root exudates, carbon subsidies to symbiotic fungi (mycorrhizae) or bacteria (e.g. nitrogen fixers), and BVOCs emissions (Figure S2). These ‘occult’ components are commonly ignored when estimating NPP, hence this bias is necessarily propagated into the *R*_a_ estimate when *R*_a_ is calculated as the difference between GPP and NPP^52^.

### Data selection

The data were obtained from more than 300 peer-reviewed articles (see also ref.^4^), adding, merging and extending previously published works worldwide on CUE or BPE ^3,8,10,22,24,53^ and covering a time-span mostly from 1995 to 2015. Data were extracted directly from the text, Tables or directly from Figures using the Unix software g3data (version 1.5.2, Jonas Frantz). In most studies, NPP, BP and GPP were estimated for the tree stand only. However, GPP estimated from CO2 flux by micrometeorological methods applies to the entire stand including ground vegetation. We therefore included only those micrometeorological studies where the forest stand was the dominant primary producer. The database contains the largest forest CUE/BPE global dataset with 244 records (197 for BPE and 47 for CUE) from >100 forest sites (including planted, managed, recently burned, N-fertilized, irrigated and CO_2_-fertilized forests; Supporting Information, Figure S1 and online Materials) representing 89 different tree species. We assume that when productivity data came from biometric (see below) measurements the reported NPP would have to be considered as actually BP because ‘occult’, non-structural and secondary carbon compounds (e.g. BVOCs or exudates) are not included. Globally, 170 records out of the total data are from temperate sites, 51 from boreal, and 23 for tropical sites, corresponding to 79 deciduous broad-leaf (DBF), 14 evergreen broad-leaf (EBF), 132 evergreen needle-leaf (ENF) and 19 mixed-forests records (MX). In some cases, multiple data sets from the same site were included, covering different years or published by different authors. From studies where data were available from more than one year, mean values across years were calculated: while we considered only those values where either NPP (or BP) and GPP referred to the same year. By using only commonly available environmental drivers to analyse the spatial variability in CUE and BPE, we were able to include almost all of the data that we found in the literature. We examined site-level effects of: average stand age (*n* = 204; range from 5 to ~500 years), mean annual temperature (MAT; *n* = 230; range –6.5 to 27.1 °C) and total annual precipitation (TAP; *n* = 232; range from ~125 to ~3500 mm yr^−1^), methods of determination (*n* = 237, see below), geographic location (latitude and longitude; *n* = 241, 64°07’ N to –42°52’ S and 155°70’ W to – 173°28’ E), elevation (*n* = 217; 5 to 2800 m, above sea level), leaf area index (LAI, *n* = 117; range from 0.4 to 13 m^2^ m^−2^), treatment (e.g.: ambient or artificially increased atmospheric CO_2_ concentration; *n* = 34), disturbance type (e.g.: fire *n* = 6; management *n* = 55), and the International Geosphere-Biosphere Programme (IGBP) vegetation classification and biomes (*n* = 244), as reported in the published articles, as potential predictors (online Materials). The methods by which GPP, NPP, BP (and Ra) were determined were included as random effects in a number of possible mixed-effects linear regression models (Table S4).

Published FPE values that were not accompanied by GPP and NPP (or by GPP and *R*_a_) or BP data (*n* = 16) were included in summary plots but excluded from the statistical analysis. Further, we excluded from statistical analysis all data where GPP and NPP were empirically determined (e.g. data obtained using fixed fractions of NPP or *R*_a_ of GPP). In just one case GPP was estimated as the sum of up-scaled *R*_a_ and NPP^54^; this study was excluded from the statistical analysis, due to small number of observations. NPP or *R*_a_ estimates obtained by process-based models (*n* = 23) were also not included in the statistical analysis (the data set lacks of BP estimations from models). No information was available on prior natural disturbance events (biotic and abiotic, e.g.: insect herbivore and pathogen outbreaks, and drought) that could in principle modify production efficiency, apart from fire. The occurrence of fire was reported by only few studies^55–57^. These data were included in the database but fire, as an explanatory factor, was not considered due to the small number of samples in which it was reported (*n* = 6).

### Estimation methods

We grouped the ‘*methods*’ into four categories:

- *Biometric*: direct tree stock measurements, or proxy data together with biomass expansion factors and the stock change as a BP component. If not otherwise stated, we assumed that the values included both above- and below-ground plant parts (*n* = 13 for GPP; *n* = 200 for NPP or BP).
- *Micromet*: micrometeorological flux measurements using the eddy-covariance technique to measure CO2 flux and partitioning methods to separate ecosystem respiration from GPP (*n* = 98 for GPP; *n* = 4 for NPP or BP).
- *Model*: model applications ranging from single mathematical equations (for canopy photosynthesis and whole-tree respiration) to more complex mechanistic process-based models to estimate GPP and *R*_a_, with NPP as the net difference between them (*n* = 53 for GPP; *n* = 24 for NPP or BP).
- *Scaling*: up-scaling of chamber-based measurements of assimilation and respiration (GPP, *Ra*) fluxes at the organ scale, or the entire stand (*n* = 73 for GPP; *n* = 9 for NPP or BP).

The difference between *‘scaling’* and *‘modelling’* lies in the data used. In the case of *‘scaling’* the data were derived from measurements at the site. ‘*Model*’ means that a dynamic process-based model was used, but with parameters calibrated and optimized at the site, based on either biometric or micrometeorological measurements.

### Data uncertainty

Uncertainties of GPP, NPP and BP data were all computed following the method based on expert judgment as described in Luyssaert et al.^52^. First, ‘gross’ uncertainty in GPP (g C m^−2^ yr^−1^) was calculated as 500 + 7.1 × (70 – |latitude|) g C m^−2^ yr^−1^ and ‘gross’ uncertainties in NPP and BP (g C m^−2^ yr^−1^) were calculated as 350 + 2.9 × (70 – |latitude|). The absolute value of uncertainty thus decreases linearly with latitude for GPP and for NPP and BP. Subsequently, as in Luyssaert et al.^52^, uncertainty was further reduced considering the methodology used to obtain each variable, by a method-specific factor (from 0 to 1), thus: final uncertainty (δ) = gross uncertainty × method-specific factor. Luyssaert et al.^52^ reports for GPP-*Micromet* a method-specific factor of 0.3 (i.e. gross uncertainty is reduced by 70% through deriving it from micrometeorological measurements), and for GPP-*Model* of 0.6, whereas for *GPP-Scaling* and *GPP-Biometric* – although not explicitly considered in Luyssaert et al.^52^ for GPP – we used the value of 0.8 and 0.3, respectively. For *BP-Biometric* and *NPP-Micromet* a reduction factor of 0.3, and for *NPP-Model* of 0.6 and *NPP-Scaling* (as obtained from chamber-based *R*_a_ measurements) of 0.8 were used. When GPP and/or NPP or BP methods were not known (*n* = 7), a factor of 1 (i.e. no reduction of uncertainty for methods used, hence maximum uncertainty) was used. The absolute uncertainties on CUE (δCUE) and BPE (δBPE) were considered as the weighted means^58^ by error propagation of each single variable (δNPP or δBP and δGPP) as follows:

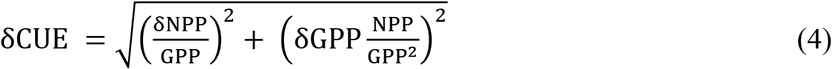

and similarly for δBPE, by substituting NPP with BP and CUE with BPE.

### Model selection

The CUE and BPE data were combined into a single variable, as sites for which both types of estimates existed did not show any significant differences between these entities (Figure 1 b). CUE values based on modelling were excluded (in our database we do not have BPE data from modelling). Tests showed that those CUE values where GPP was estimated with micrometeorological methods was systematically higher, compared to values based on biometric or scaling methods. Only data with complete information on CUE, MAT, age, TAP, and latitude were used. This resulted to a data selection corresponding to 142 observations.

First, to use the information as completely as possible, a full additive model was constructed (equation 1 in the main text). The method used for estimation of GPP (*GPPmeth*) was specified as a random effect on the intercept as visual inspection of the data suggested that CUE values were smaller where *‘scaling’* was used to estimate GPP compared to cases where ‘ *micromet’* was used to estimate GPP.

In equation (1) the variable ‘age’ represents the development status of the vegetation, i.e. either average age of the canopy forming trees or the period since the last major disturbance, while the other three parameters represent different aspects of the climate. The absolute latitude, |lat|, was chosen as a proxy of radiation climate, i.e. day length and the seasonality of daily radiation. The term *ηZ_GPPmeth_* represents the random effect on the intercept due to the different methods of estimating GPP.

These variables were not independent (Table S1). If the different driver variables contain information that is not included in any of the other driver variables, multiple linear regression is nonetheless able separate the individual effects. If, on the contrary, two variables mainly exert the same effect on the response variable (CUE) this can be seen in an ANOVA based model comparison. These considerations led us to the selection procedure: starting with the full model (equation 1), and then comparing it with all possible reduced models (Table S2). The result of this analysis is the model with the smallest number of parameters that does not significantly differ from the full model.

We also examined, whether there were any significant interactions of predictor variables, which was not the case.

We used the `R` function lmer from the `R` package lme4^59^ to fit the mixed and ordinary multiple linear models to the data. The model residuals were tested for normality (Anderson Darling test of non-normality, `R` function and test of the `R` package nortest^60^). For models that did not take a random intercept regarding *‘GPPmeth’* into account (16 to 30 in Table S2) the Anderson Darling test found significant deviation from normality of the model residuals, these models were excluded from the analysis. The remaining models were compared with each other using the function ANOVA of the `R` package lmerTest^61^. This resulted in a 15 · 15 matrix of model comparisons in which the full model turned out to be significantly different from all other models. This shows that it was not possible to reduce the model any further.

The same analysis was also performed with a log-transformed version of equation (1):

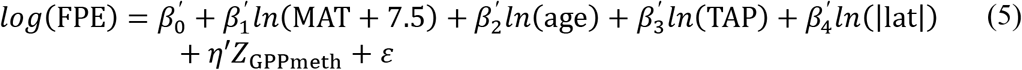

where 7.5 °C was added to MAT in order to make its minimum 1 °C. Note that the linear model from the log-transformed variables differs from the untransformed linear model. The coefficients, here noted with a prime, can be interpreted on the basis of the back-transformed model. Contrary to the untransformed linear model where effects are additive, the back-transformed model is a multiplicative effect model, with the slope parameters as exponents for each variable and the intercept (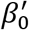 as power of *e*). As with the untransformed model, negative slope parameter values lower CUE, positive increase it with increasing driver variable values.

The results from this analysis was, as with the original additive model (equation 1), (*i*) the full model could not be reduced any further and (*ii*) although now multiplicative, the directions of the effects were exactly the same as with the additive model, i.e. the predicted CUE increased with increasing MAT, TAP and |lat| but decreased with increasing age.

The AIC and BIC values were lower for the log-transformed model compared to the untransformed model, with AIC values of –169.7 and –157.2 for the log-transformed and untransformed models, respectively, and BIC values of –149.0 and –136.5 for the log-transformed and untransformed models, respectively. The coefficients and model performance parameters of the untransformed and the log-transformed models are shown in Table 1 and Table S4. The adjusted squared correlation coefficient were similar, i.e. 0.306 for the untransformed and 0.321 for the log-transformed model. Despite considerable uncertainty of the CUE values, it was possible to derive significant, systematic, linear relationships between the four driver variables and CUE or ln (CUE). Both model variants show, in principle, the same direction and magnitudes of the effects. These results qualify the log-transformed model only a little better than the linear model. It can be concluded that CUE (or ln (CUE)) from a global data set of a large variety of forests is significantly positively affected by MAT, TAP and |lat|, and significantly negatively affected by age.

Because the parameters of the untransformed, additive model are much easier to interpret, we use the additive model in the main text and use the log-transformed model only as a confirmation of trends found in the additive model.

### Outputs from TRENDY v.7

We used the simulations from eight Dynamic Global Vegetation Models (DGVMs) performed in the framework of the TRENDY v.7 project^2,62^ (http://dgvm.ceh.ac.uk/node/9; data downloaded 27 November 2018). Models that did not provide NPP and GPP at plant functional type level were excluded because of the need of having CUE in pure forests without significant contributions from shrubs, grassland or crops. The selection comprises the following models: ISAM, JULES, LPJ-GUESS, CABLE-POP, ORCHIDEE, ORCHIDE-CNP, JSBACH and SDGVM (for references on models see ref.^2,62^ and Table S5). All the models represent the surface fluxes of CO_2_, water and the dynamics of carbon pools in response to changes in climate, atmospheric CO2 concentration, and land use change across a global grid. However, processes underlying the exchanges of water and carbon are based on different formulations among models.

In the TRENDY protocol all DGVMs were forced with common historical climate fields and atmospheric CO2 concentrations over the period from 1700 to 2017. Climate fields were taken from the CRU-JRA55 dataset^2^, whereas the time series of atmospheric CO_2_ concentrations were derived from the combination of ice core records and atmospheric observations. Land-use change was taken into account in the used simulations S3. However, similar simulations but without land use (S2) were also tested showing no differences. CUE was estimated as NPP/GPP (where NPP is commonly obtained in models by removing *R*_a_ from GPP) for the forest plant functional types simulated to be present in each grid cell. BP was not, conversely, directly available – but neither indirectly obtainable – by model outputs. The model outputs refer to 1995-2015 for comparability with the data analysis.

## Acknowledgments

The authors thanks R.H. Waring, S. Vicca, M. Campioli, and E. Grieco for early constructive comments and thoughtful suggestions; S. Noce for the map of data points. We thank efforts from all site investigators and their funding agencies;

## Author contributions

A.C. A.I. and I.C.P. conceived the paper, A.C., A.S., A.I., A.C., R.A. analysed data, A.C, A.I., A.C., R.A., M.F.-M., and I.C.P. wrote the manuscript. All authors contributed substantially to discussions and revisions;

## Data and materials availability

All data supporting this study are available in the supplementary materials and will be publicly available at Figshare repository upon acceptance (doi:10.6084/m9.figshare.11917908). Correspondence and requests for materials should be addressed to A.I..

## References

1. Reich, P. B. et al. Boreal and temperate trees show strong acclimation of respiration to warming. Nature 531, 633–636 (2016).

2. Le Quéré, C. et al. Global Carbon Budget 2018. Earth Syst. Sci. Data 10, 2141–2194 (2018).

3. Vicca, S. et al. Fertile forests produce biomass more efficiently. Ecol. Lett. 15, 520–526 (2012).

4. Collalti, A. & Prentice, I. C. Is NPP proportional to GPP? Waring’s hypothesis 20 years on. Tree Physiol. 39, 1473–1483 (2019).

5. Waring, R. H., Landsberg, J. J. & Williams, M. Net primary production of forests: a constant fraction of gross primary production? Tree Physiol. 18, 129–134 (1998).

6. Cannell, M. G. R. & Thornley, J. H. M. Modelling the Components of Plant Respiration: Some Guiding Principles. Ann. Bot. 85, 45–54 (2000).

7. Collalti, A. et al. Plant respiration: Controlled by photosynthesis or biomass? Glob. Chang. Biol. In press (2019) doi:10.1111/gcb.14857.

8. Campioli, M. et al. Biomass production efficiency controlled by management in temperate and boreal ecosystems. Nat. Geosci. 8, 843–846 (2015).

9. Fernández-Martínez, M. et al. Nutrient availability as the key regulator of global forest carbon balance. Nat. Clim. Chang. 4, 471–476 (2014).

10. DeLucia, E. H., Drake, J. E., Thomas, R. B. & Gonzalez-meler, M. A. Forest carbon use efficiency: is respiration a constant fraction of gross primary production? Glob. Chang. Biol. 13, 1157–1167 (2007).

11. He, Y. et al. Global vegetation biomass production efficiency constrained by models and observations. Glob. Chang. Biol. 26, 1474–1484 (2020).

12. Medlyn, B. E. & Dewar, R. C. Comment on the article by R. H. Waring, J. J. Landsberg and M. Williams relating net primar production to gross primary production. Tree Physiol. 19, 137–138 (1999).

13. van Dam, N. M. & Bouwmeester, H. J. Metabolomics in the Rhizosphere: Tapping into Belowground Chemical Communication. Trends Plant Sci. 21, 256–265 (2016).

14. Heinemeyer, A. et al. Exploring the ‘overflow tap’ theory: linking forest soil CO2 fluxes and individual mycorrhizosphere components to photosynthesis. Biogeosciences 9, 79–95 (2012).

15. Preece, C., Farré-Armengol, G., Llusià, J. & Peñuelas, J. Thirsty tree roots exude more carbon. Tree Physiol. 38, 690–695 (2018).

16. Guenther, A. The contribution of reactive carbon emissions from vegetation to the carbon balance of terrestrial ecosystems. Chemosphere 49, 837–844 (2002).

17. Kuhn, U. et al. Strong correlation between isoprene emission and gross photosynthetic capacity during leaf phenology of the tropical tree species Hymenaea courbaril with fundamental changes in volatile organic compounds emission composition during early leaf development. Plant. Cell Environ. 27, 1469–1485 (2004).

18. Amthor, J. S. The McCree–de Wit–Penning de Vries–Thornley Respiration Paradigms: 30 Years Later. Ann. Bot. 86, 1–20 (2000).

19. Medlyn, B. E. et al. Effects of elevated [CO2] on photosynthesis in European forest species: a meta-analysis of model parameters. Plant. Cell Environ. 22, 1475–1495 (1999).

20. Drake, J. E., Davis, S. C., Raetz, L. M. & DeLucia, E. H. Mechanisms of age-related changes in forest production: the influence of physiological and successional changes. Glob. Chang. Biol. 17, 1522–1535 (2011).

21. Collalti, A. et al. The sensitivity of the forest carbon budget shifts across processes along with stand development and climate change. Ecol. Appl. 29, (2019).

22. Litton, C. M., Raich, J. W. & Ryan, M. G. Carbon allocation in forest ecosystems. Glob. Chang. Biol. 13, 2089–2109 (2007).

23. Piao, S. et al. Forest annual carbon cost: a global-scale analysis of autotrophic respiration. Ecology 91, 652–661 (2010).

24. Tang, J., Luyssaert, S., Richardson, A. D., Kutsch, W. & Janssens, I. A. Steeper declines in forest photosynthesis than respiration explain age-driven decreases in forest growth. Proc. Natl. Acad. Sci. 111, 8856–8860 (2014).

25. Ryan, M. G., Phillips, N. & Bond, B. J. The hydraulic limitation hypothesis revisited. Plant, Cell Environ. 29, 367–381 (2006).

26. Reich, P. B. et al. Scaling of respiration to nitrogen in leaves, stems and roots of higher land plants. Ecol. Lett. 11, 793–801 (2008).

27. Mori, S. et al. Mixed-power scaling of whole-plant respiration from seedlings to giant trees. Proc. Natl. Acad. Sci. 107, 1447–1451 (2010).

28. Johnson, D. W. Progressive N limitation in forests: review and implications for long-term responses to elevated CO2. Ecology 87, 64–75 (2006).

29. Gill, A. L. & Finzi, A. C. Belowground carbon flux links biogeochemical cycles and resource-use efficiency at the global scale. Ecol. Lett. 19, 1419–1428 (2016).

30. Way, D. A. & Sage, R. F. Elevated growth temperatures reduce the carbon gain of black spruce [Picea mariana (Mill.) B.S.P.]. Glob. Chang. Biol. 14, 624–636 (2008).

31. Collalti, A. et al. Thinning Can Reduce Losses in Carbon Use Efficiency and Carbon Stocks in Managed Forests Under Warmer Climate. J. Adv. Model. Earth Syst. 10, 2427–2452 (2018).

32. Michaletz, S. T., Cheng, D., Kerkhoff, A. J. & Enquist, B. J. Convergence of terrestrial plant production across global climate gradients. Nature 512, 39–43 (2014).

33. Malhi, Y. The productivity, metabolism and carbon cycle of tropical forest vegetation. J. Ecol. 100, 65–75 (2012).

34. Merganičová, K. et al. Forest carbon allocation modelling under climate change. Tree Physiol. (2019) doi:10.1093/treephys/tpz105.

35. Sala, A. & Hoch, G. Height-related growth declines in ponderosa pine are not due to carbon limitation. Plant. Cell Environ. 32, 22–30 (2009).

36. Dietze, M. C. et al. Nonstructural Carbon in Woody Plants. Annu. Rev. Plant Biol. 65, 667–687 (2014).

37. Metcalfe, D. B. et al. Shifts in plant respiration and carbon use efficiency at a large-scale drought experiment in the eastern Amazon. New Phytol. 187, 608–621 (2010).

38. Ibrom, A. et al. Variation in photosynthetic light-use efficiency in a mountainous tropical rain forest in Indonesia. Tree Physiol. 28, 499–508 (2008).

39. Larcher, W. Physiological Plant Ecology. (Springer-Verlag Berlin Heidelberg, 2003).

40. Drake, J. E. et al. Does physiological acclimation to climate warming stabilize the ratio of canopy respiration to photosynthesis? New Phytol. 211, 850–863 (2016).

41. Gifford, R. M. Plant respiration in productivity models: conceptualisation, representation and issues for global terrestrial carbon-cycle research. Funct. Plant Biol. 30, 171–186 (2003).

42. O’Leary, B. M., Asao, S., Millar, A. H. & Atkin, O. K. Core principles which explain variation in respiration across biological scales. New Phytol. 222, 670–686 (2019).

43. Smith, N. G. & Dukes, J. S. Plant respiration and photosynthesis in global-scale models: incorporating acclimation to temperature and CO2. Glob. Chang. Biol. 19, 45–63 (2013).

44. Wang, H. et al. Acclimation of leaf respiration consistent with optimal photosynthetic capacity. Glob. Chang. Biol. n/a, (2020).

45. He, Y., Piao, S., Li, X., Chen, A. & Qin, D. Global patterns of vegetation carbon use efficiency and their climate drivers deduced from MODIS satellite data and process-based models. Agric. For. Meteorol. 256–257, 150–158 (2018).

46. Griffin, K. L. & Prager, C. M. Where does the carbon go? Thermal acclimation of respiration and increased photosynthesis in trees at the temperate-boreal ecotone. Tree Physiol. 37, 281–284 (2017).

47. O’sullivan, O. S. et al. Thermal limits of leaf metabolism across biomes. Glob. Chang. Biol. 23, 209–223 (2017).

48. Sperling, O., Earles, J. M., Secchi, F., Godfrey, J. & Zwieniecki, M. A. Frost Induces Respiration and Accelerates Carbon Depletion in Trees. PLoS One 10, e0144124–e0144124 (2015).

49. Thornley, J. H. M. Plant growth and respiration re-visited: maintenance respiration defined – it is an emergent property of, not a separate process within, the system – and why the respiration: photosynthesis ratio is conservative. Ann. Bot. 108, 1365–1380 (2011).

50. Landsberg, J. J., Waring, R. H. & Williams, M. Commentary on the assessment of NPP/GPP ratio. Tree Physiol. (2020) doi:10.1093/treephys/tpaa016.

51. Chapin, F. S. et al. Reconciling Carbon-cycle Concepts, Terminology, and Methods. Ecosystems 9, 1041–1050 (2006).

52. Luyssaert, S. et al. CO 2 balance of boreal, temperate, and tropical forests derived from a global database. Glob. Chang. Biol. 13, 2509–2537 (2007).

53. Campioli, M. et al. Evaluating the convergence between eddy-covariance and biometric methods for assessing carbon budgets of forests. Nat. Commun. 7, 13717 (2016).

54. Curtis, P. S. et al. Respiratory carbon losses and the carbon-use efficiency of a northern hardwood forest, 1999–2003. New Phytol. 167, 437–456 (2005).

55. Law, B. E., Thornton, P. E., Irvine, J., Anthoni, P. M. & Van Tuyl, S. Carbon storage and fluxes in ponderosa pine forests at different developmental stages. Glob. Chang. Biol. 7, 755–777 (2001).

56. Dore, S. et al. Carbon and water fluxes from ponderosa pine forests disturbed by wildfire and thinning. Ecol. Appl. 20, 663–683 (2010).

57. Goulden, M. L. et al. Patterns of NPP, GPP, respiration, and NEP during boreal forest succession. Glob. Chang. Biol. 17, 855–871 (2011).

58. Slob, W. Uncertainty Analysis in Multiplicative Models. Risk Anal. 14, 571–576 (1994).

59. Bates, D., Mächler, M., Bolker, B. & Walker, S. Fitting Linear Mixed-Effects Models Using lme4. J. Stat. Software; Vol 1, Issue 1 (2015).

60. Gross, J. & Ligges, U. nortest: Tests for Normality. R package version 1.0-4. (2015).

61. Kuznetsova, A., Brockhoff, P. B. & Christensen, R. H. B. lmerTest Package: Tests in Linear Mixed Effects Models. J. Stat. Software; Vol 1, Issue 13 (2017).

62. Sitch, S. et al. Recent trends and drivers of regional sources and sinks of carbon dioxide. Biogeosciences 12, 653–679 (2015).

