## Supplementary material for "Forest production efficiency increases with growth temperature": Supplemetary Materials

### Supplementary Information

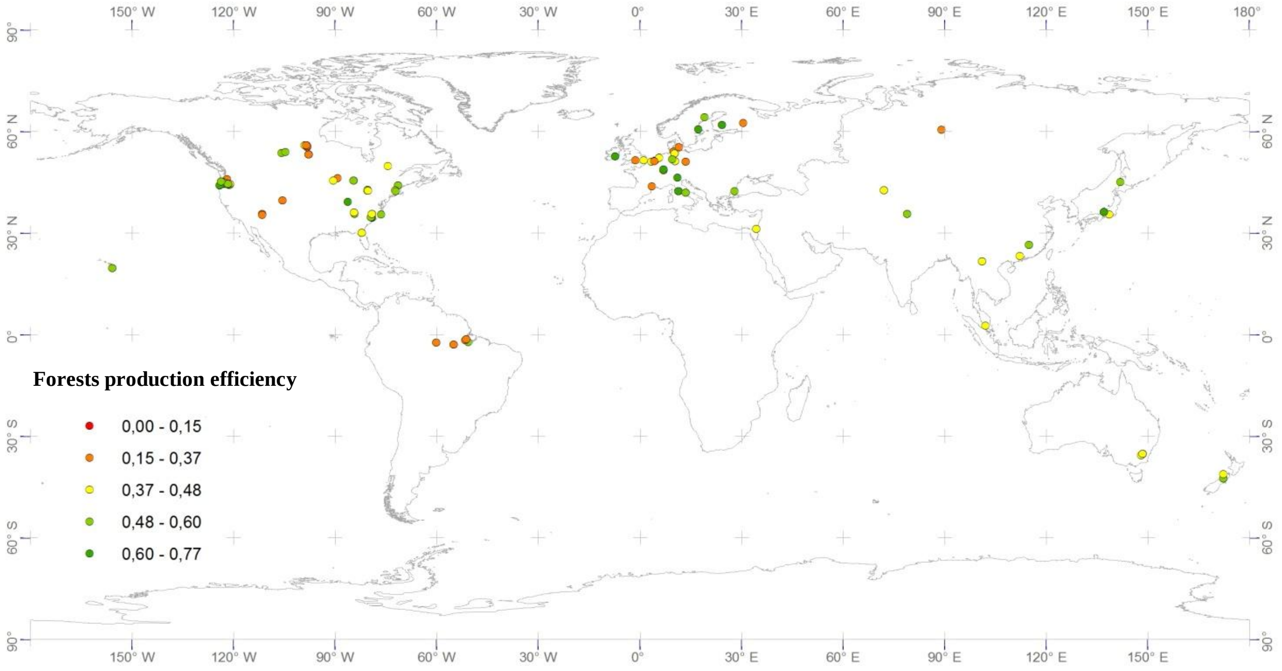


**Figure S1 |** A global map of forest sites used to create a database of carbon use and biomass production efficiency (grouped as 'Forest production efficiency' in the figure legend)

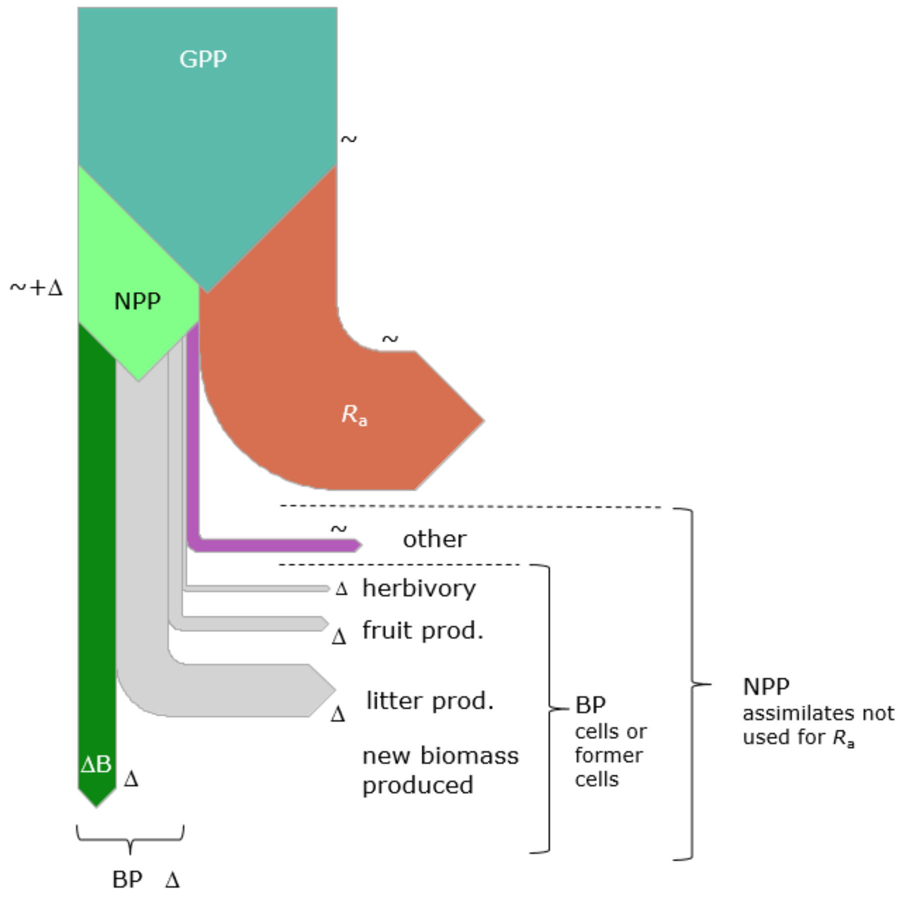


**Figure S2 |** Schematic of carbon flows in and through plants, and their relationships to the quantities defined in eq. (3). Measurements are by sequential inventory (Δ) or gas and solute exchange (~) methods.

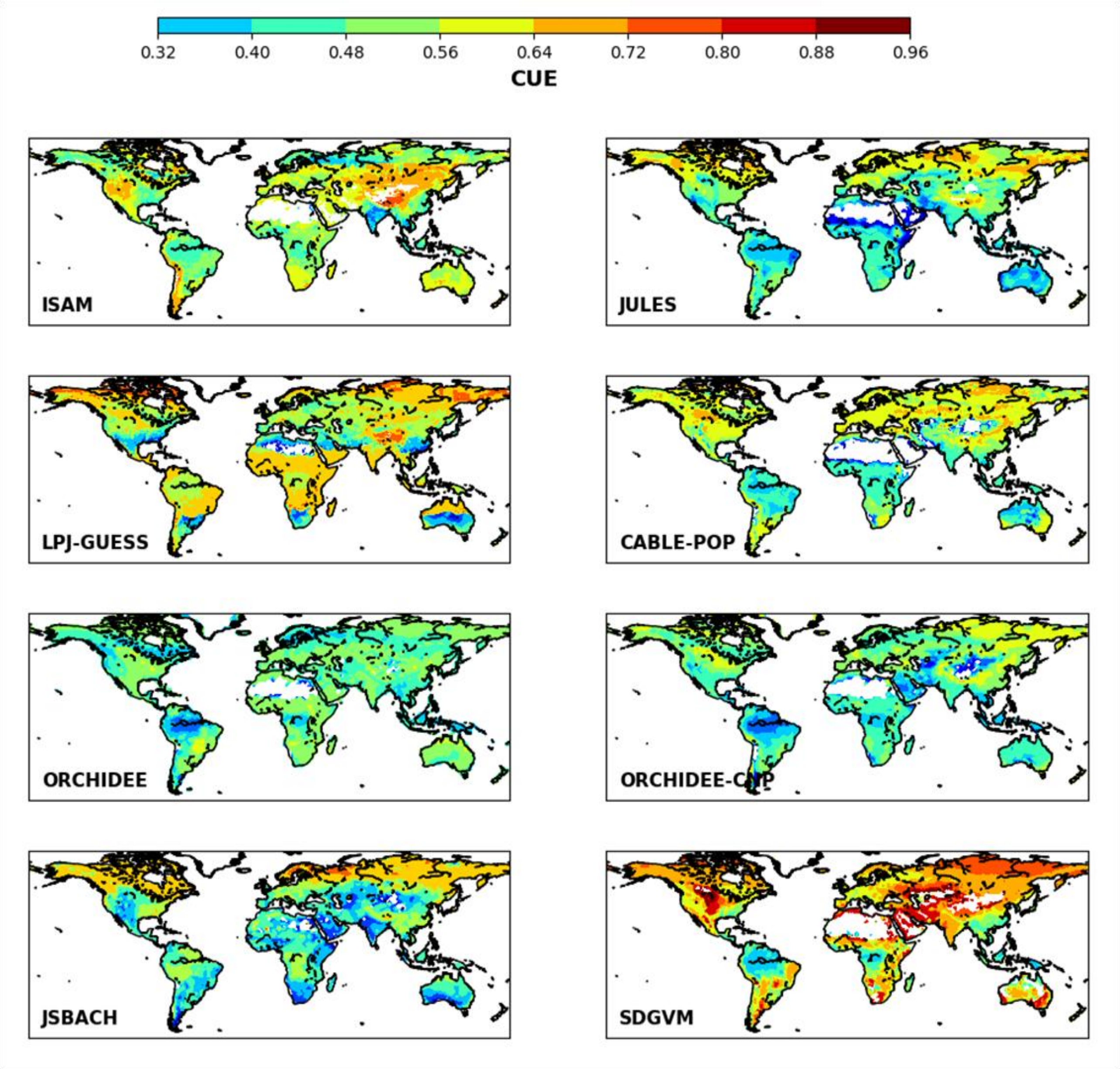


**Figure S3 |** Global patterns of vegetation carbon use efficiency (CUE) derived from TRENDY v.7 process-based models: ISAM, JULES, LPJ-GUESS, CABLE-POP, ORCHIDEE, ORCHIDEE-CNP, JSBACH and SDGVM, averaged from 1995 to 2015.

**Table S1 |** Pearson's correlation matrix of the model driver variables.

| | **CUE** | **age** | **MAT** | **TAP** | **\|lat\|** |
|---|---|---|---|---|---|
| **CUE** | | –0.178 | 0.234 | 0.354 | –0.242 |
| **age** | n.s. | | –0.868 | –0.555 | 0.721 |
| **MAT** | n.s. | *** | | 0.506 | –0.843 |
| **TAP** | n.s. | * | * | | –0.773 |
| **\|lat\|** | n.s. | ** | *** | *** | |

**Table S2 |** Fixed and random intercept variables of the models examined in step 1. The 'x' in the model matrix below represents the terms in equation (1) that include the variable of the respective column header. The intercept ($\beta_0$) is part of all models (not included in the mode matrix).

| No. | MAT | age | TAP | \|latitude\| | random intercept GPP method |
|---|---|---|---|---|---|
| 1 | x | x | x | x | x |
| 2 | x | x | x | | x |
| 3 | x | x | | x | x |
| 4 | x | | x | x | x |
| 5 | | x | x | x | x |
| 6 | x | x | | | x |
| 7 | x | | x | | x |
| 8 | x | | | x | x |
| 9 | | x | x | | x |
| 10 | | x | | x | x |
| 11 | | | x | x | x |
| 12 | x | | | | x |
| 13 | | x | | | x |
| 14 | | | x | | x |
| 15 | | | | x | x |
| 16 | x | x | x | x | |
| 17 | x | x | x | | |
| 18 | x | x | | x | |
| 19 | x | | x | x | |
| 20 | | x | x | x | |
| 21 | x | x | | | |
| 22 | x | | x | | |
| 23 | x | | | x | |
| 24 | | x | x | | |
| 25 | | x | | x | |
| 26 | | | x | x | |
| 27 | x | | | | |
| 28 | | x | | | |
| 29 | | | x | | |
| 30 | | | | x | |

**Table S3** | Site years with both CUE and BPE estimates

| Site ID | Location | Species | Age |
|---|---|---|---|
| 1 | Bartlett Experimental Forest | *Acer saccharum, Fagus grandifolia, Fraxinus americar* | 80 |
| 2 | Bornhoved Lake Beech, Germany | *Fagus sylvatica* | 111 |
| 3 | Dooary forest | *Picea sitchensis* | 18 |
| 4 | Harvard forest | *Quercus rubra, Acer rubrum, Taxus canadensis* | 100 |
| 5 | Hesse, France | *Fagus sylvatica* | 32 |
| 6 | Hesse, France | *Fagus sylvatica* | 40 |
| 7 | Hyytiälä | *Pinus sylvestris* | 47 |
| 8 | Norunda | *Scots pine , Norway spruce* | 105 |
| 9 | Oregon Transect Ecosystem Research - Metolius River Valley | *Pinus ponderosa* | 148 |
| 10 | Prince Albert Canada | *Picea mariana* | 115 |
| 11 | Prince Albert Canada | *Pinus banksiana* | 63 |
| 12 | Prince Albert Canada | *Populus tremuloides* | 68 |
| 13 | Takayama, Japan | *Betula ermanii, B. platyphylla, Quercus mongolia* | 40 |

**Table S4 |** Model performance parameters for the full *log-transformed* model, i.e. equation (5).

| | Estimate | Std Error | df | t-value | p-value | significance |
|---|---|---|---|---|---|---|
| **Intercept** | | | | | | |
| **'Micromet'** | –0.57 | | | | | |
| **'scaling'** | –0.65 | | | | | |
| **Intercept ($\boldsymbol{\beta}'_0$)** | –0.61302 | 0.108603 | 32.5 | –5.6446 | 2.9E–06 | *** |
| **MAT ($\boldsymbol{\beta}'_1$)** | 0.006129 | 0.002409 | 132.0 | 2.544659 | 0.012089 | * |
| **Age ($\boldsymbol{\beta}'_2$)** | –0.00042 | 0.000127 | 132.3 | –3.28225 | 0.001317 | ** |
| **TAP ($\boldsymbol{\beta}'_3$)** | 6.51E–05 | 2.25E–05 | 132.2 | 2.894311 | 0.004446 | ** |
| **absLat ($\boldsymbol{\beta}'_4$)** | 0.003737 | 0.001631 | 132.6 | 2.291606 | 0.023505 | * |

**Table S5 |** Description of autotrophic respiration ($R_a$) and its components, growth ($R_g$) and maintenance ($R_m$) respiration, and reserves (non-structural carbon pool, NSC) for the eight TRENDY v.7 models used in the data vs. model comparison. For general definition of Acclimation and Adaption see Smith *&* Dukes (2013).

| | Model | | | | | | | |
|---|---|---|---|---|---|---|---|---|
| **Process** | **ISAM** | **JULES** | **LPJ-GUESS** | **CABLE-POP** | **ORCHIDEE** | **ORCHIDEE-CNP** | **JSBACH** | **SDGVM** |
| **GPP** | Farquhar | Collatz | Haxeltine & Prentice | Farquhar, extended to account for co-ordination of rate-determining steps in photosynthesis. | Farquhar | Farquhar | Farquhar | Farquhar |
| **Growth Respiration*** (growth respiration coefficient, $r_G$) | Not available | 25% | 33% | >15%: magnitude depends on leaf P/N ratio | 28% | 28% | 25% | 25% |
| **Maintenance Respiration** (Temperature dependence) | Fixed $Q_{10}$ | Fixed $Q_{10}$. Bell-shaped function with peak rates at 32°C | Constant modified Arrhenius function in response to temperature | Variable $Q_{10}$ as in Atkin *et al.* (2016) | Constant modified Arrhenius function in response to temperature | Constant modified Arrhenius function in response to temperature | Constant exponential response to temperature plus high-temperature inhibition | Stem and root: exponential increase, leaf (day): exponential increase capped at 30 °C, leaf (night): none |
| **Maintenance Respiration** | Not available | Depends on biomass N | Depends on biomass N | Depends on biomass N | Depends on biomass N | Depends on biomass N | Linear dependence on | Stem and root: proportion of |

| | | | | | | | | |
|---|---|---|---|---|---|---|---|---|
| (Biomass dependence) | | concentration | concentration | concentration | concentration | concentration | leaf area index, but LAI not depending on biomass | live biomass**, leaf (day): proportion of leaf N*, leaf (night): proportion of leaf N |
| **Acclimation of $R_a$** | Not available | Previous 10 days temperature (only for leaves) | No | Previous three months temperature (for leaves, stems and fine roots) | No | No | No | No |
| **Adaptation of $R_a$** | Not available | No | No | No | No | No | No | No |
| **Autotrophic Respiration** | Sum of $R_g$ + $R_m$ | Sum of $R_g$ + $R_m$ | Sum of $R_g$ + $R_m$ | Sum of $R_g$ + $R_m$ | Sum of $R_g$ + $R_m$ | Sum of $R_g$ + $R_m$ | Sum of $R_g$ + $R_m$ | Sum of $R_g$ + $R_m$ |
| **NPP** | NPP = (GPP–$R_m$) (1–$r_G$) | NPP = (GPP–$R_m$) (1–$r_G$) | NPP = (GPP–$R_m$) (1–$r_G$) | NPP = (GPP–$R_m$) (1–$r_G$) | NPP = (GPP–$R_m$) (1–$r_G$) | NPP = (GPP–$R_m$) (1–$r_G$) | NPP = (GPP–$R_m$) (1–$r_G$) | NPP = (GPP–$R_m$) (1–$r_G$) |
| **Reserves** | Not available | No | No | No | Yes | Yes | Yes | Yes |
| ***References*** | King *et al.* (1996) ; Kheshgi & Jain 2003 | Atkin *et al.* (2008); Huntingford *et al*. (2017) | Smith *et al.* (2014) | Haverd *et al.* (2018) | Krinner *et al.* (2005) | Goll *et al.* (2017) | Raddatz *et al.*, (2007); Mauritsen *et al.* (2019) | Woodward *et al*. (1995) |

* $R_g$ = (GPP-$R_m$) * $r_G$

** but also multiplied by soil water limitation scalar

### References

Atkin O. K. *et al*. (2008). Using temperature-dependent changes in leaf scaling relationships to quantitatively account for thermal acclimation of respiration in a coupled global climate–vegetation model. *Global Change Biology*, 14, 2709–2726, doi: 10.1111/j.1365-2486.2008.01664.x

Atkin, O. K. *et al*. (2016). Global variability in leaf respiration in relation to climate, plant functional types and leaf traits. *New Phytologist*, 211, 1142–1142.

Huntingford C. *et al*. (2017). Implications of improved representations of plant respiration in a changing climate. *Nature Communications*, doi: 10.1038/s41467-017-01774-z

Kheshgi, H. S., and A. K. Jain, (2003) Projecting future climate change: Implications of carbon cycle model inter-comparisons, *Global Biogeochem. Cycles*, 17(2), 1047, doi:10.1029/2001GB001842

King A. W. *et al*. (1995). In search of the missing carbon sink: a model of terrestrial biospheric response to land-use change and atmospheric $CO_2$, *Tellus B: Chemical and Physical Meteorology*, 47:4, 501-519, DOI:10.3402/tellusb.v47i4.16064

Krinner G. *et al*. (2005). A dynamic global vegetation model for studies of the coupled atmosphere-biosphere system. *Global Biogeochemical Cycles*, doi:10.1029/2003GB002199

Goll D.S. *et al*. (2017). A representation of the phosphorus cycle for ORCHIDEE (revision 4520). *Geosci. Model Dev*.,

Haverd V. *et al*. (2018). A new version of the CABLE land surface model (Subversion revision r4601) incorporating land use and land cover change, woody vegetation demography, and a novel optimisation-based approach to plant coordination of photosynthesis. *Geosci. Model Dev*., 11, 2995–3026, https://doi.org/10.5194/gmd-11-2995-201

Mauritsen T. *et al*. (2019). Developments in the MPI-M Earth System Model version 1.2 (MPI-ESM1.2) and Its Response to Increasing $CO_2$, *Journal of Advances in Modeling Earth System*, 11, https://doi.org/10.1029/2018MS001400

Raddatz T. J., *et al*. (2007), Will the tropical land biosphere dominate the climate-carbon cycle feedback during the twenty-first century?, *Clim. Dyn*.,29, 565–574

Smith B. *et al*. (2014). Implications of incorporating N cycling and N limitations on primary production in an individual-based dynamic vegetation model. *Biogeosciences*, doi:10.5194/bg-11-2027-2014

Woodward F.I. *et al*. (1995). A global land primary productivity and phyto-geography model, *Global Biogeochemical Cycles*, https://doi.org/10.1029/95GB02432
